## Supplemental Info for "Helical reconstruction of VP39 reveals principles for baculovirus nucleocapsid assembly"

**Supplementary Fig. 1: Data processing scheme.**

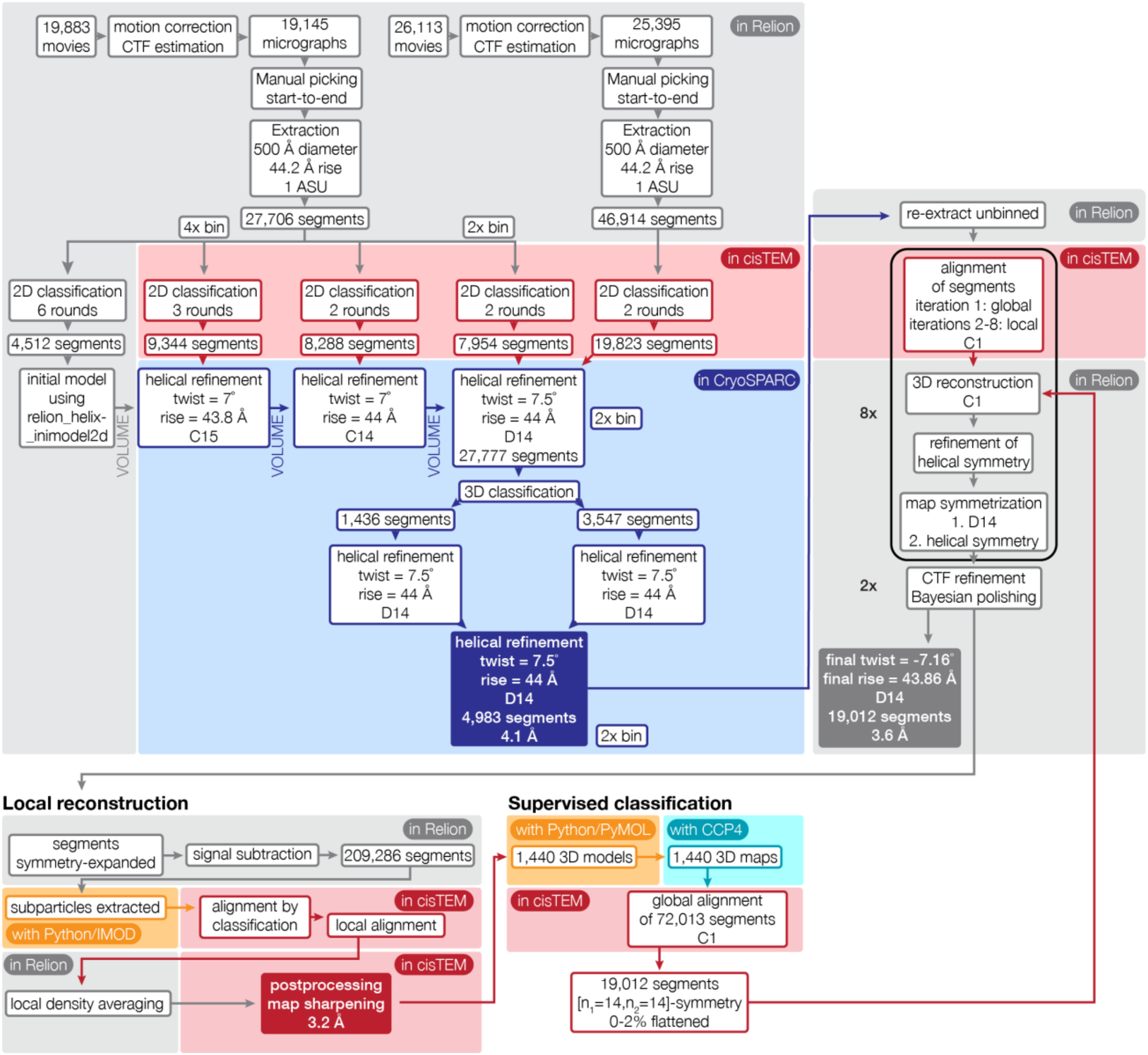

**Supplementary Fig. 2: Determination of helical symmetry parameters by Fourier-Bessel indexing.**

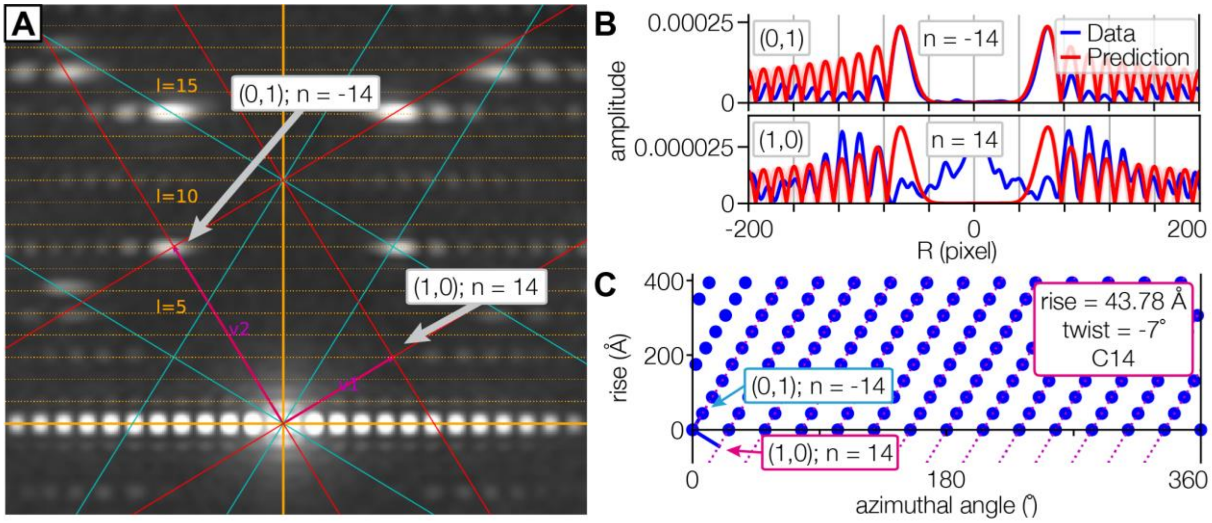


(A) Average power spectrum of all segments of the final [$n_1$=14, $n_2$=14] reconstruction. Layer lines are shown as dashed lines in orange. Unit vectors $\mathbf{v_1}$ and $\mathbf{v_2}$ (pink) were used to generate a 2D lattice (red) consistent with peaks in the power spectrum. The mirror lattice is depicted in cyan.

(B) Amplitude diagrams for measured (blue) and predicted (red) data on layer line l=8 (for vector (0,1)) and on layer line l=3 (for vector (1,0)) of the power spectrum in (A) for Bessel order n=14.

(C) Real-space 2D lattice calculated from the Fourier-space lattice in (A) with the corresponding helical symmetry parameters.

**Supplementary Fig. 3: Alignment of baculoviral VP39 sequences.**

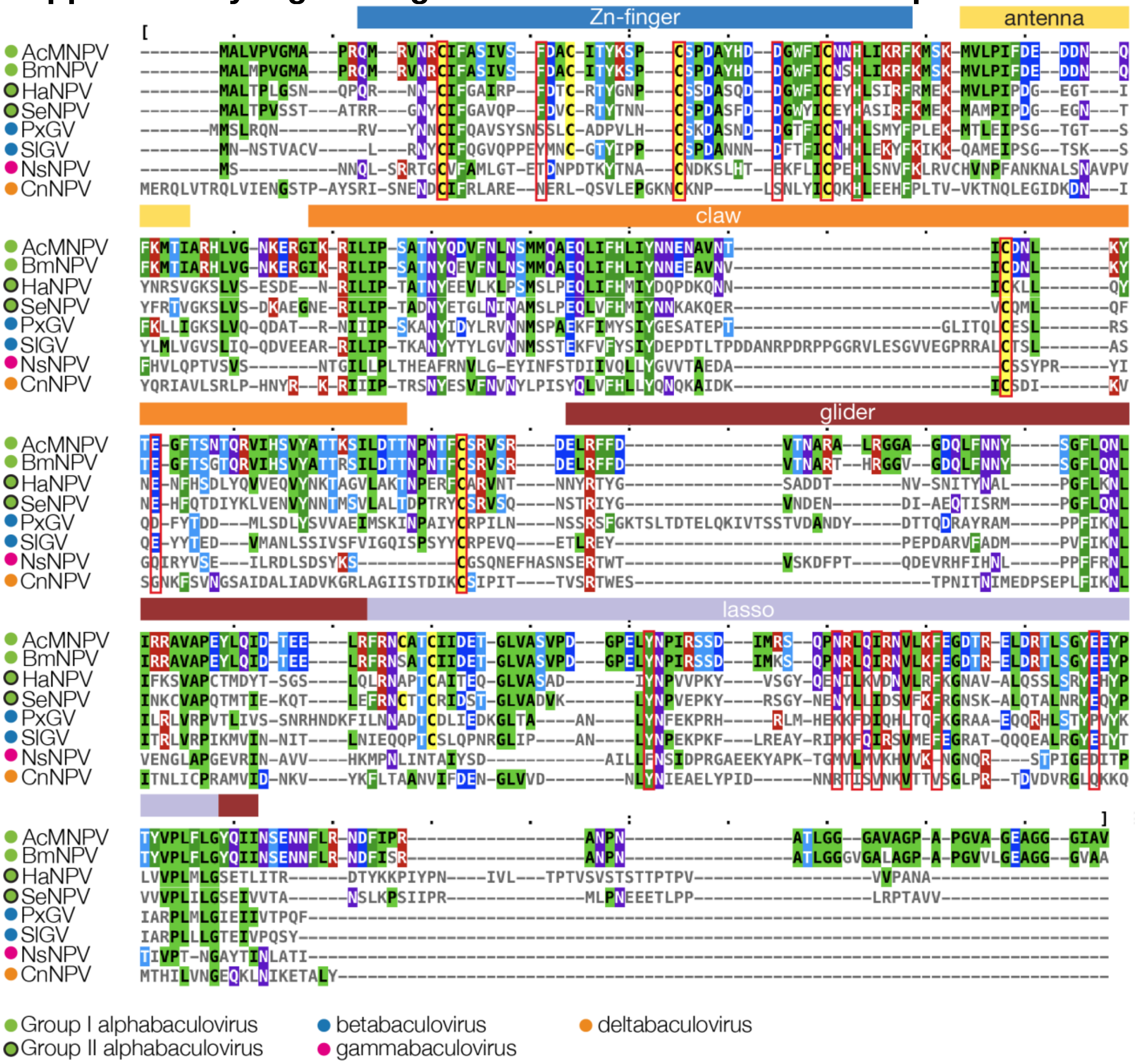


8 selected VP39 sequences are shown from an alignment of 73 sequences. Residues are colored by identity and conserved residues listed in Supplementary Table 4 are boxed in red. Sequence alignment was performed using MAFFT and visualized with MView.[1,2] AcMNPV: *A. californica* MNPV; BmNPV: *Bombyx mori* NPV; HaNPV: *H. armigera* NPV; SeNPV: *S. exigua* NPV; PxGV: *P. xylostella* GV; SlGV: *S. litura* GV; NsNPV: *N. sertifer* NPV; CnNPV: *C. nigripalpus* NPV.

**Supplementary Fig. 4: Phylogenetic tree of baculoviral VP39 calculated from 73 sequences.**

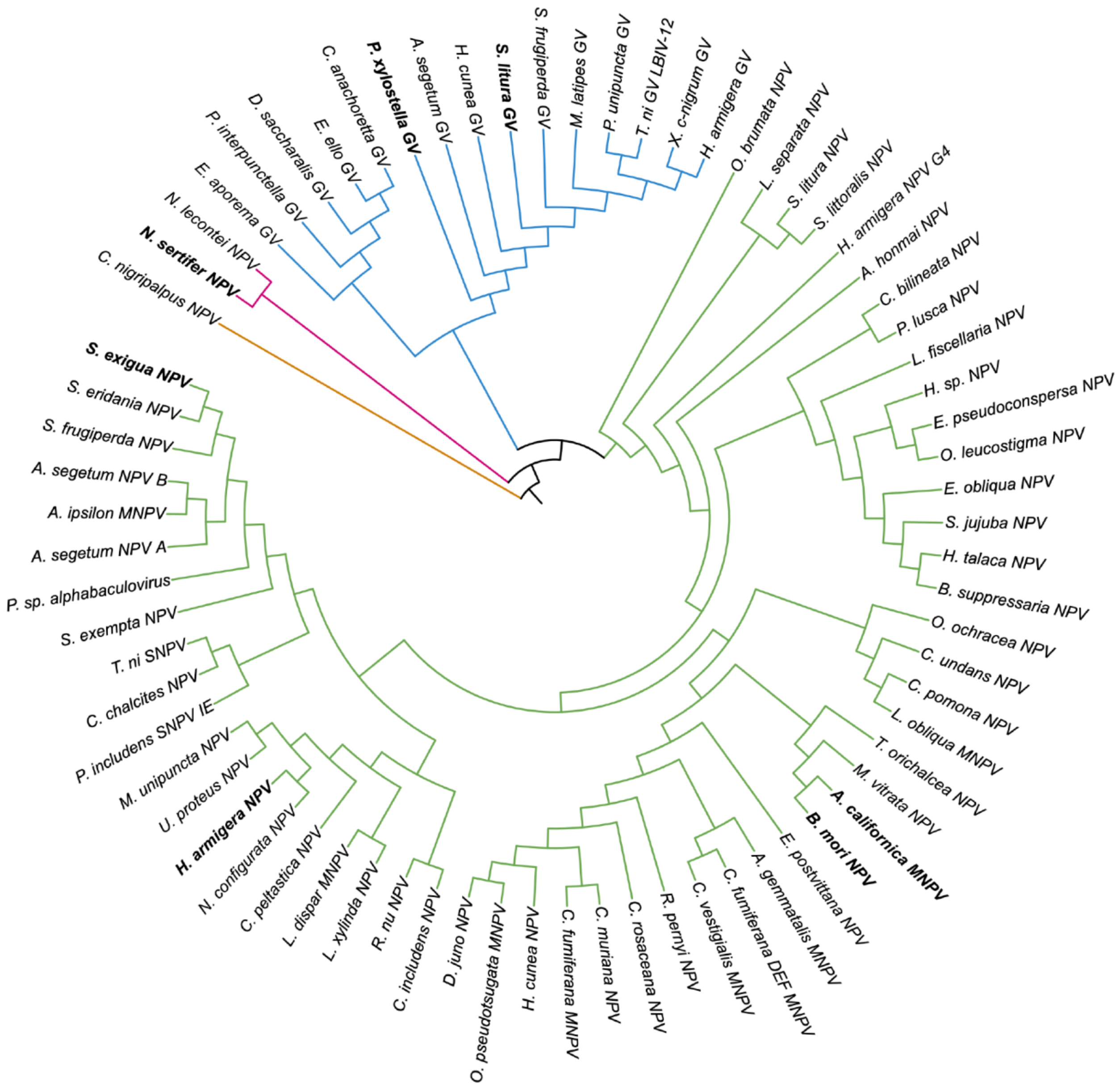


● alphabaculovirus ● betabaculovirus ● gammabaculovirus ● deltabaculovirus

Viral species sequences that were used for helical reconstruction (AcMNPV) and for structure predictions are highlighted in bold. Branches are colored according to viral genera: alphabaculovirus (green), betabaculovirus (blue), gammabaculovirus (pink), deltabaculovirus (orange). The phylogenetic tree was calculated with Jalview[3] using the Neighbor Joining algorithm with a BLOSUM62 substitution matrix[4] and visualized using the Interactive Tree of Life iTOL v. 6.7.4[5]. Branch lengths are ignored in the tree representation.

**Supplementary Fig. 5: Confidence depiction for baculoviral VP39 dimer model predictions by AlphaFold2.**

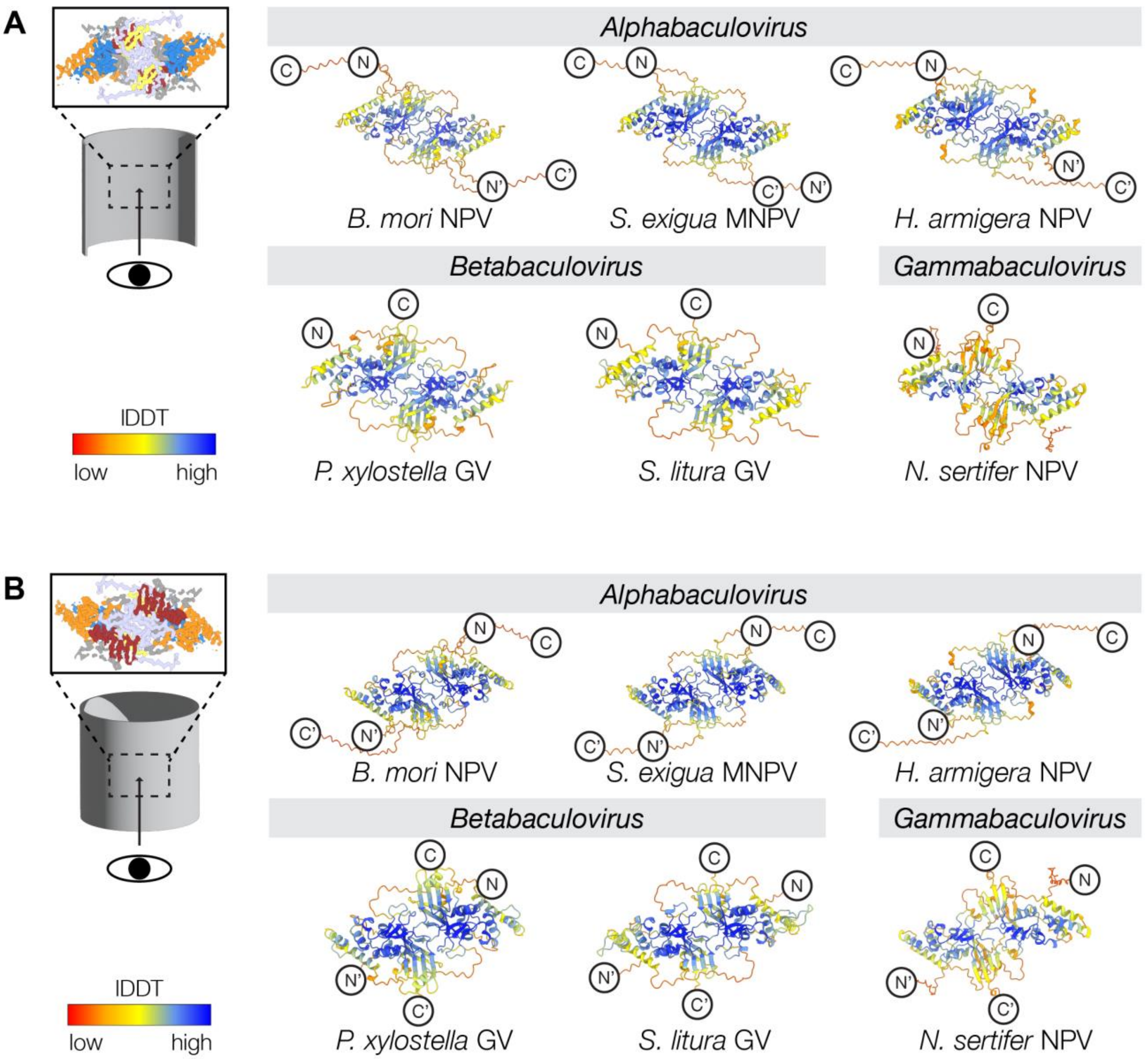


(A) Luminal side of baculoviral predicted VP39 dimer models. The structures are colored by prediction confidence from low (red) to high (blue), as indicated by the lDDT color bar. Prediction confidence is expressed through an lDDT-score, which is assigned a color[6]. Notably, prediction for the extended termini and inter-dimer stretches of the lasso loop are low.

(B) Exterior side of baculoviral VP39 dimer models, colored by prediction confidence.

NPV: nucleopolyhedrovirus; GV: granulovirus; lDDT: local Distance Difference Test.

**Supplementary Fig. 6: Comparison of observed and calculated power spectra of the five most populated, non-flattened classes after supervised classification.**

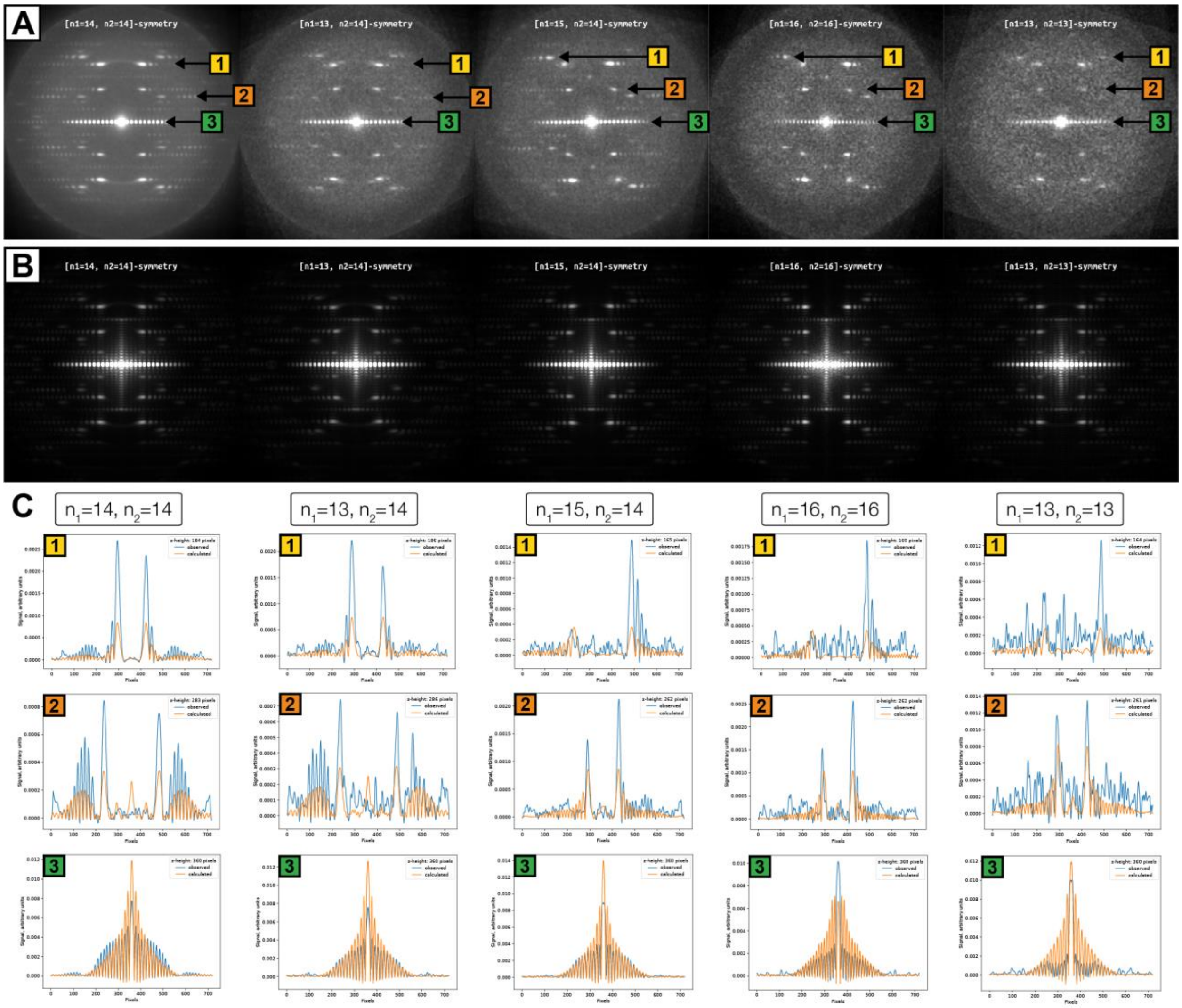


(A) Averaged power spectra of segments, which classified to a 3D reference with the indicated symmetry. Numbered arrows specify layer lines that were selected for direct comparison with the corresponding layer lines of calculated power spectra (shown in B) in panel C. Segments were rotated according to the in-plane angle from the 3D alignment before calculating the power spectrum.

(B) Averaged power spectra calculated from 3D reference projections after orienting the 3D references according to the alignment parameters as observed for each segment.

Projections were rotated according to the in-plane angle from the 3D alignment before calculating the power spectrum.

(C) Layer line profiles from power spectra of segments (shown in panel A; “observed”; blue lines) and calculated power spectra from reference projections (shown in panel B; “calculated”; orange lines) for prominent layer lines (denoted as 1, yellow; 2, red; 3, green in panel A) for each helical symmetry. For better comparison, layer line profiles were baseline corrected and arbitrarily scaled (linear scale factor). One pixel corresponds to a spatial frequency of 0.000166 $Å^{-1}$.

**Supplementary Fig. 7: Classification analysis of the subparticle stack used for local reconstruction.**

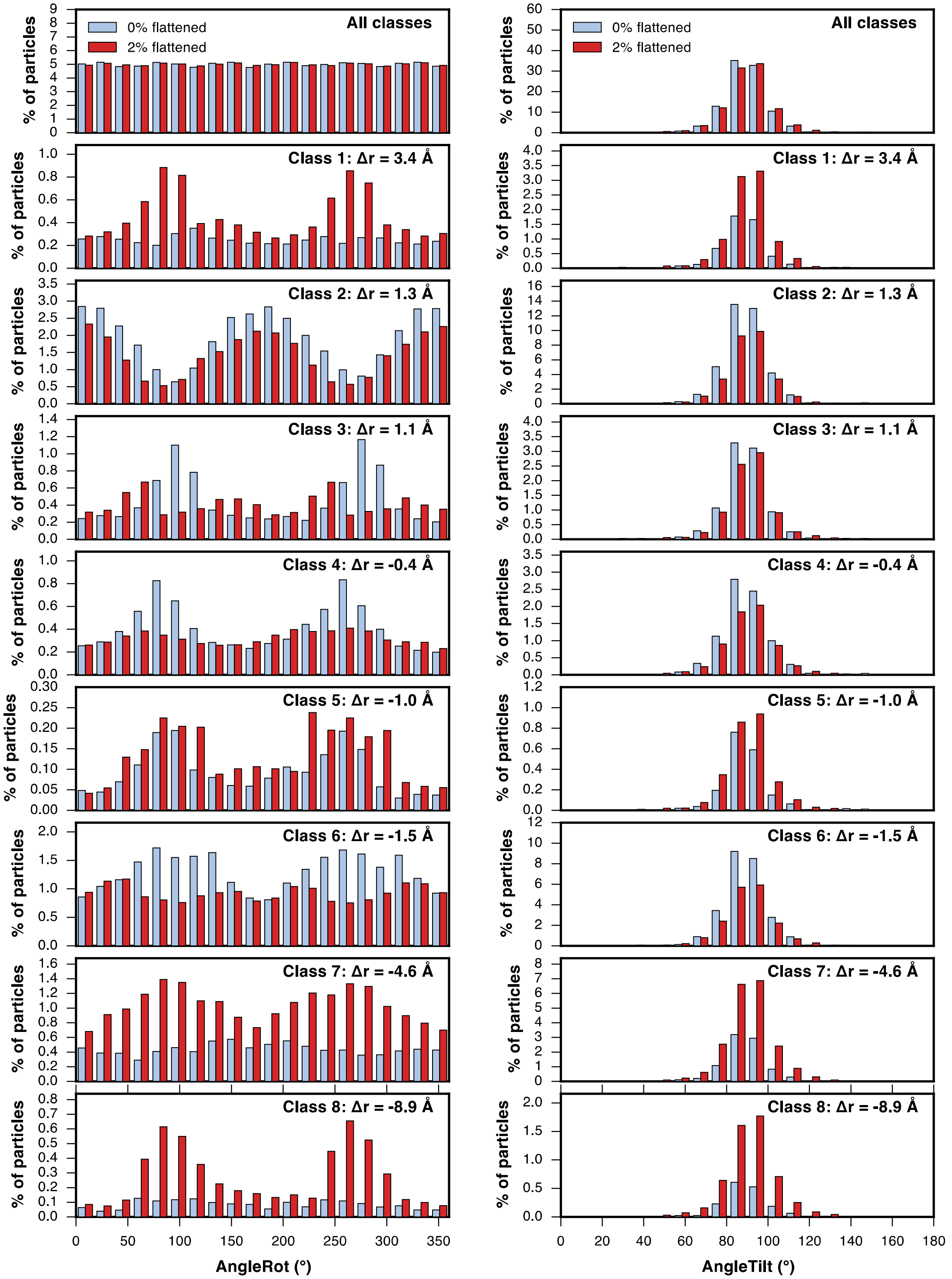

A subparticle stack extracted from 19,012 segments ([$n_1$=14, $n_2$=14] symmetry and from non-flattened up to 2% flattened as determined by supervised classification) was classified into eight classes without changing the alignment. The plots show the distribution of the alignment angles (AngleRot, rotation around the helical axis; AngleTilt, out-of-plane rotation) for particles extracted from non-flattened (blue bars) and 2% flattened (red bars) segments, respectively. The top row shows the angles for particles from all classes, the following rows show each class individually with decreasing radial shifts from the helical axis ($\Delta$r). Classes with the largest radial shifts from the helical axis (classes 1, 7 and 8) show a substantial bias towards particles derived from 2% flattened segments.

**Supplementary Fig. 8: Reconstructions of the three most populated classes after supervised classification.**

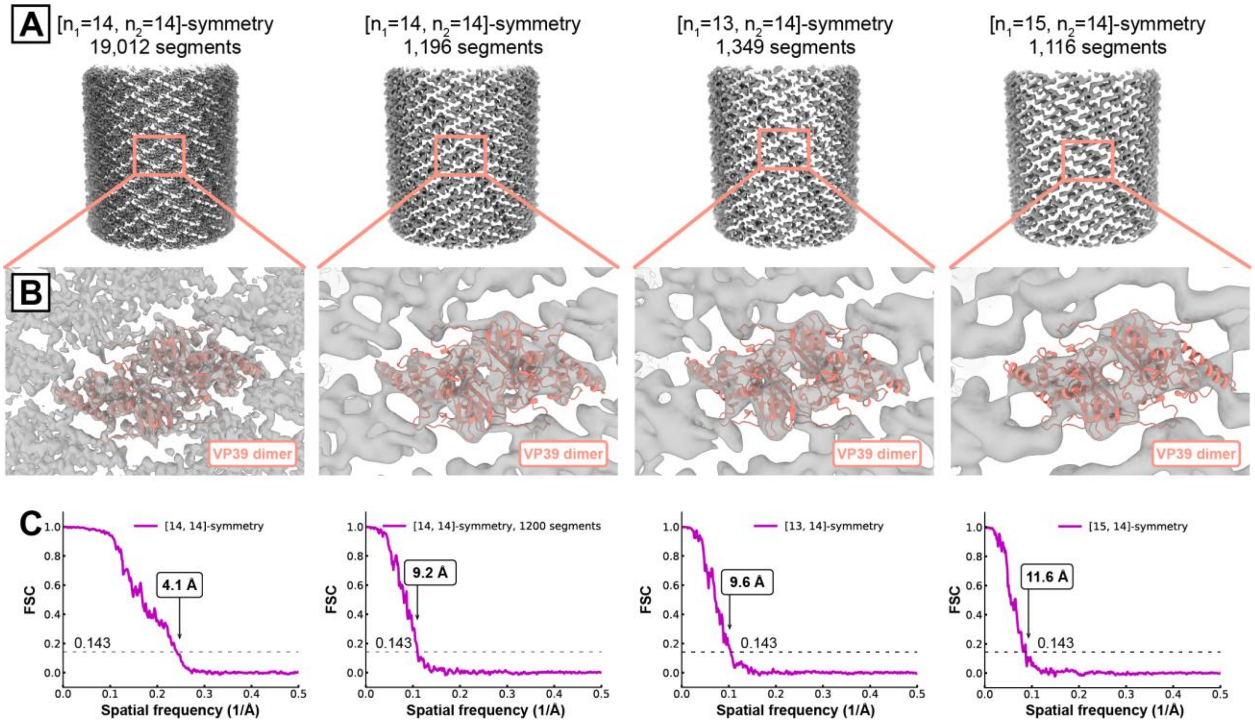


(A) Reconstructed volumes for segments, which classified to a 3D reference of 0-2% flattening with the indicated symmetry. Left: reconstruction of all 19,012 segments with [$n_1$=14, $n_2$=14] symmetry at 4.1 Å using the simplified reconstruction protocol (see Methods). Center left: reconstructed volume of 1,196 randomly selected segments with [$n_1$=14, $n_2$=14] symmetry at 9.2 Å using the simplified reconstruction protocol to compare to reconstructions of helical symmetries with fewer segments. Center right and right: reconstructed volumes of segments, which classified to a 3D reference of [$n_1$=13, $n_2$=14] and [$n_1$=15, $n_2$=14] symmetry, respectively, at 9.6 Å and 11.6 Å (see Supplementary Table 5 for reconstruction statistics). Contrastingly to the main reconstruction protocol, only one cycle of the following steps was performed instead of two cycles: segment alignment in cisTEM, reconstruction in C1, map symmetrization, and relion_ctf_refine. The main reconstruction protocol includes an additional helical symmetry search, map symmetrization in D14 and relion_motion_refine, which this simplified reconstruction protocol does not include. In the simplified reconstruction protocol applied here, map symmetrization is first performed using local symmetry (for protomers within a z_percentage of 0.3), followed by helical symmetry, whereas map symmetrization in the main reconstruction protocol is first performed using D14 rotational symmetry, followed by helical symmetry.

(B) Close-up of dimer region with the VP39 dimer fitted into the map.

(C) FSC curves calculated from half maps for each reconstruction.

### Supplementary Table 1: Cryo-EM data collection and model statistics.

| | AcMNPV VP39 nucleocapsid |
|---|---|
| **Data collection** | |
| Electron microscope | Titan Krios |
| Magnification | 60606 |
| Voltage (kV) | 300 |
| Defocus range (µm)[a] | 0.4–1.9 |
| Pixel size (Å) | 0.825 |
| Number of movies | 45984 |
| **Helical reconstruction, unbinned** | |
| Number of images | 19,012 |
| Box size (pixels) | 912 |
| Symmetry imposed | $[n_1=14, n_2=14]$ [b], D14 followed by helical symmetry |
| Helical twist (˚) | -7.16 |
| Helical rise (Å) | 43.86 |
| Map resolution (Å)[c] | 3.6 |
| **Local reconstruction** | |
| Number of images | 209,286 |
| Box size (pixels) | 912 |
| Symmetry imposed | C1 |
| Map resolution (Å)[c] | 3.2 |
| **Model statistics** | |
| EMD accession identifier | EMD-41133 |
| PDB accession identifier | 8TAF |
| Refinement resolution (Å) | 3.2 |
| CC (mask) | 0.78 |
| Model composition | |
| Non-hydrogen atoms | 39440 |
| Protein residues | 2464 |
| Ligands | Zn: 8 |
| *B* factors | |
| Protein ($Å^2$) | 49.77 (mean) |
| Ligand ($Å^2$) | 86.97 (mean) |
| R.m.s deviations | |
| Bond lengths (Å) | 0.003 |
| Bond angles (˚) | 0.519 |
| Validation | |
| MolProbity score | 1.43 |
| Clash score | 3.12 |
| Romater outliers (%) | 0.4 |
| Ramachandran plot | |

| | |
|---|---|
| Favored (%) | 95.34 |
| Allowed (%) | 4.49 |
| Outliers (%) | 0.16 |

[a] Approximate range of underfocus.

[b] [$n_1$=14, $n_2$=14] symmetry was applied by reconstruction in C1, then applying D14 symmetry followed by applying helical symmetry.

[c] Resolution where FSC between masked half maps drops below 0.143.

#### Supplementary Table 2: Analysis of the AcMNPV VP39 dimer interface.

Interface surface area and interaction types within the AcMNPV VP39 dimer were analyzed using the PDBePISA webserver[7].

| Interface 1 | Interface 2 | Area [$Å^2$] | Hydrogen bonds | Salt bridges | $\Delta G_{sol}$ |
|---|---|---|---|---|---|
| Chain O | Chain J | 3548.3 | 23 | 3 | -35.5 |

**List of hydrogen bonds comprising the AcMNPV VP39 dimer interface**

| Chain O | Chain J | Distance [Å] |
|---|---|---|
| Asp 44 [OD2] | Tyr 288 [HZ1] | 1.53 |
| Val 63 [H] | Glu 289 [O] | 2.06 |
| Val 63 [O] | Tyr 291 [H] | 1.84 |
| Phe 67 [O] | Tyr 250 [HH] | 1.70 |
| Asp 68 [OD2] | Phe 274 [H] | 2.22 |
| Asn 72 [HD21] | Phe 274 [O] | 2.24 |
| Arg 225 [HH11] | Pro 247 [O] | 2.08 |
| Arg 225 [O] | Tyr 250 [H] | 1.87 |
| Arg 227 [HH12] | Tyr 291 [OH] | 1.94 |
| Asp 245 [O] | Arg 225 [HH12] | 2.04 |
| Pro 247 [O] | Arg 225 [HH11] | 1.98 |
| Tyr 250 [H] | Arg 225 [O] | 2.12 |
| Tyr 250 [HH] | Phe 67 [O] | 1.75 |
| Arg 254 [HE] | Asp 70 [O] | 1.79 |
| Arg 254 [HH21] | Asp 70 [O] | 1.97 |
| Arg 254 [HH22] | Gln 73 [OE1] | 2.23 |
| Leu 272 [H] | Glu 69 [OE2] | 2.40 |
| Lys 273 [HZ2] | Asp 70 [OD2] | 2.40 |
| Phe 274 [H] | Asp 68 [OD2] | 1.72 |
| Phe 274 [O] | Asn 72 [HD21] | 1.68 |
| Tyr 288 [HH] | Asp 44 [OD2] | 1.65 |
| Glu 289 [O] | Val 63 [H] | 2.32 |
| Tyr 291 [H] | Val 63 [O] | 1.79 |

**List of salt bridges comprising the AcMNPV VP39 dimer interface**

| Chain O | Chain J | Distance [Å] |
|---|---|---|
| Lys 75 [NZ] | Asp 282 [OD1] | 3.74 |
| Arg 225 [NZ] | Asp 245 [OD2] | 3.50 |
| Lys 273 [NZ] | Asp 70 [OD2] | 3.05 |

#### Supplementary Table 3: Analysis of the VP39 helical repeat unit contacts.

Interface surface area and interaction types between VP39 helical repeat units were analysed using the PDBePISA webserver[7].

##### LATERAL INTERACTIONS

| Interface 1 | Interface 2 | Area [Å$^2$] | Hydrogen bonds | Salt bridges | $\Delta G_{sol}$ |
|---|---|---|---|---|---|
| Chain M | Chain J | 605.8 | 1 | 0 | -12.9 |
| Chain P | Chain R | 614.3 | 1 | 0 | -14.6 |
| | *Average:* | *610.0* | | | *-13.8* |

##### List of hydrogen bonds at the lateral interface

| Chain M | Chain J | Distance [Å] |
|---|---|---|
| Gln 146 | Cys 169 | 2.10 |

| Chain P | Chain R | Distance [Å] |
|---|---|---|
| Gln 146 | Cys 169 | 1.87 |

##### TYPE-II AXIAL INTERACTIONS

| Interface 1 | Interface 2 | Area [Å$^2$] | Hydrogen bonds | Salt bridges | $\Delta G_{sol}$ |
|---|---|---|---|---|---|
| Chain P | Chain O | 703.0 | 10 | 2 | -4.0 |
| Chain M | Chain L | 714.9 | 11 | 2 | -8.5 |
| | *Average:* | *709.0* | | | *-6.2* |

**List of hydrogen bonds at the type-II axial interface**

| Chain P | Chain O | Distance [Å] |
|---|---|---|
| Pro 263 [O] | Asp 277 [H] | 2.25 |
| Gln 267 [O] | Glu 275 [H] | 1.95 |
| Gln 267 [H] | Glu 275 [O] | 2.36 |
| Arg 269 [H] | Lys 273 [O] | 2.50 |
| Arg 269 [HE] | Glu 275 [OE2] | 2.45 |
| Lys 273 [H] | Arg 269 [O] | 2.43 |
| Lys 273 [O] | Arg 269 [H] | 2.49 |
| Glu 275 [H] | Gln 267 [O] | 2.07 |
| Glu 275 [O] | Gln 267 [H] | 2.48 |
| Asp 277 [H] | Pro 263 [O] | 2.00 |

| Chain M | Chain L | Distance [Å] |
|---|---|---|
| Pro 263 [O] | Asp 277 [H] | 2.02 |
| Gln 267 [O] | Glu 275 [H] | 2.44 |
| Gln 267 [H] | Glu 275 [O] | 2.45 |
| Arg 269 [O] | Lys 273 [H] | 2.39 |
| Val 271 [O] | Val 271 [H] | 2.25 |
| Val 271 [H] | Val 271 [O] | 2.18 |
| Lys 273 [H] | Arg 269 [O] | 2.46 |
| Lys 273 [O] | Arg 269 [H] | 2.10 |
| Glu 275 [H] | Gln 267 [O] | 2.30 |
| Glu 275 [OE2] | Arg 269 [HE] | 2.16 |
| Asp 277 [H] | Pro 263 [O] | 2.37 |

**List of salt bonds at the type-II axial interface**

| Chain P | Chain O | Distance [Å] |
|---|---|---|
| Arg 269 [NE] | Glu 275 [OE2] | 3.12 |
| Glu 275 [OE1] | Arg 269 [NE] | 3.12 |

| Chain M | Chain L | Distance [Å] |
|---|---|---|
| Arg 269 [NE] | Glu 275 [OE2] | 3.86 |
| Glu 275 [OE2] | Arg 269 [NE] | 2.76 |

#### TYPE-I AXIAL INTERACTIONS

| Interface 1 | Interface 2 | Area [Å$^2$] | Hydrogen bonds | Salt bridges | $\Delta G_{sol}$ |
|---|---|---|---|---|---|
| Chain Q | Chain O | 439.1 | 4 | 0 | -4.2 |
| Chain P | Chain J | 449.3 | 5 | 0 | -3.3 |
| Chain R | Chain M | 450.4 | 4 | 0 | -2.9 |
| Chain N | Chain L | 440.7 | 5 | 0 | -3.1 |
| | *Average:* | *444.9* | | | *-3.4* |

**List of hydrogen bonds at the type-I axial interface**

| Chain Q | Chain O | Distance [Å] |
|---|---|---|
| Ser 306 [O] | Arg 265 [HE] | 2.28 |
| Asn 308 [OD1] | Leu 266 [H] | 2.21 |
| Asn 308 [HD22] | Leu 266 [O] | 2.27 |
| Leu 311 [H] | Asn 264 [O] | 2.35 |

| Chain P | Chain J | Distance [Å] |
|---|---|---|
| Ser 306 [O] | Arg 265 [HE] | 1.64 |
| Ser 306 [O] | Arg 265 [HH21] | 2.14 |
| Glu 307 [OE1] | Asn 106 [HD22] | 1.83 |
| Asn 308 [OD1] | Leu 266 [H] | 2.36 |
| Leu 311 [H] | Asn 264 [O] | 2.32 |

| Chain R | Chain M | Distance [Å] |
|---|---|---|
| Ser 306 [O] | Arg 265 [HE] | 1.91 |
| Ser 306 [O] | Arg 265 [HH21] | 2.16 |
| Glu 307 [OE2] | Asn 106 [HD22] | 2.04 |
| Leu 311 [H] | Asn 264 [O] | 2.43 |

| Chain N | Chain L | Distance [Å] |
|---|---|---|
| Ser 306 [O] | Arg 265 [HE] | 1.94 |
| Ser 306 [O] | Arg 265 [HH21] | 2.19 |
| Glu 307 [OE2] | Asn 106 [HD22] | 1.88 |
| Asn 308 [OD1] | Leu 266 [H] | 2.48 |
| Leu 311 [H] | Asn 264 [O] | 2.45 |

#### AXIAL-LATERAL INTERACTIONS

| Interface 1 | Interface 2 | Area [Å$^2$] | Hydrogen bonds | Salt bridges | $\Delta G_{sol}$ |
|---|---|---|---|---|---|
| Chain P | Chain M | 538.4 | 1 | 0 | -5.0 |

**List of hydrogen bonds at the axial-lateral interface**

| Chain P | Chain M | Distance [Å] |
|---|---|---|
| Glu 139 [OE2] | Phe 26 [H] | 2.49 |

#### Supplementary Table 4: List of sequence-conserved residues of VP39

73 VP39 sequences across baculoviruses (55 alphabaculoviruses, 15 betabaculoviruses, 2 gammabaculoviruses and 1 deltabaculovirus) were aligned using MAFFT[1] (see methods and Supplementary Fig. 3).

| Residue[a] | Role |
|---|---|
| Cys 18 | $Zn^{2+}$ coordination |
| Phe 26[b] | Axial-lateral inter-subunit contact |
| Cys 36 | $Zn^{2+}$ coordination |
| Asp 44[c] | Intra-dimer contact |
| Cys 49 | $Zn^{2+}$ coordination |
| His 52 | $Zn^{2+}$ coordination |
| Cys 132 | Candidate disulfide partner[d] |
| Glu 139[c,e] | Axial-lateral inter-subunit contact |
| Cys 169 | Lateral inter-subunit contact, candidate disulfide partner[d] |
| Tyr 250[f] | Intra-dimer contact |
| Asn 264[b] | Type-I axial inter-subunit contact |
| Leu 266[b] | Type-I axial inter-subunit contact |
| Ile 268 | unknown |
| Val 271 | Type-II axial inter-subunit contact |
| Phe 274[c] | Intra-dimer contact |
| Glu 289[g] | Intra-dimer contact |

[a]AcMNPV VP39 residue numbering

[b]conserved in alphabaculoviruses only

[c]conserved in alpha- and betabaculoviruses only

[d]Extended Data Fig. 7

[e]present as a glutamate or an aspartate

[f]Phe in gammabaculovirus

[g]Asp in gammabaculovirus

**Supplementary Table 5: Statistics for reconstructions from segments after supervised classification for helical symmetries of the five most populated classes.**

| Symmetry | Number of segments | cisTEM score[b] | Resolution[c] | Resolution[d] |
|---|---|---|---|---|
| [14, 14] | 19012 | 18.1 | 7.9 | 4.1 |
| [14, 14][a] | 1196 | 18.1 | 13.4 | 9.2 |
| [13, 14] | 1349 | 15.4 | 14.2 | 9.6 |
| [15, 14] | 1116 | 13.3 | 18.4 | 11.6 |
| [16, 16] | 204 | 11.0 | 37.6 | N. A. |
| [13, 13] | 224 | 10.8 | 62.7 | N. A. |

[a] 1,196 segments randomly selected from the 19,012 segments.

[b] Average alignment score after supervised classification.

[c] Resolution after reconstruction in C1 without symmetrization using alignment parameters from supervised classification.

[d] Resolution after alignment and relion_ctf_refine with symmetrization.

#### References


1. Katoh, K., Rozewicki, J. & Yamada, K.D. MAFFT online service: multiple sequence alignment, interactive sequence choice and visualization. *Briefings in Bioinformatics* **20**, 1160-1166 (2017).
2. Madeira, F. et al. Search and sequence analysis tools services from EMBL-EBI in 2022. *Nucleic acids research* **50**, W276-W279 (2022).
3. Waterhouse, A.M., Procter, J.B., Martin, D.M.A., Clamp, M. & Barton, G.J. Jalview Version 2—a multiple sequence alignment editor and analysis workbench. *Bioinformatics* **25**, 1189-1191 (2009).
4. Saitou, N. & Nei, M. The neighbor-joining method: a new method for reconstructing phylogenetic trees. *Mol Biol Evol* **4**, 406-25 (1987).
5. Letunic, I. & Bork, P. Interactive Tree Of Life (iTOL) v5: an online tool for phylogenetic tree display and annotation. *Nucleic Acids Research* **49**, W293-W296 (2021).
6. Mariani, V., Biasini, M., Barbato, A. & Schwede, T. lDDT: a local superposition-free score for comparing protein structures and models using distance difference tests. *Bioinformatics* **29**, 2722-8 (2013).
7. Krissinel, E. & Henrick, K. Inference of macromolecular assemblies from crystalline state. *J Mol Biol* **372**, 774-97 (2007).
