## Extended Data for "Helical reconstruction of VP39 reveals principles for baculovirus nucleocapsid assembly"

### Extended Data Fig. 1: Variation in tube diameter.

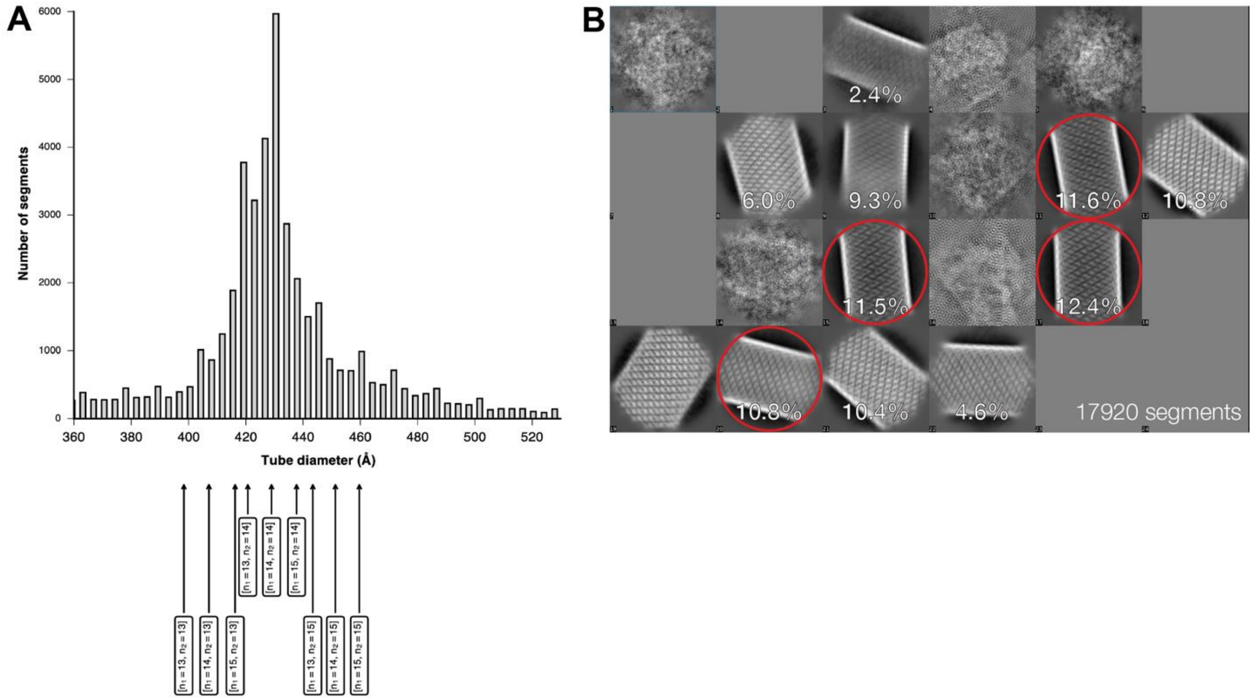


(A) Distribution of segment widths of 72,013 extracted segments. Segment widths were measured by aligning the helical axis based on the zero layer line signal of the power spectrum, projecting the density on a line and then measuring the distance between the two top peaks. Histogram bins with tube diameters <360 Å or >530 Å are not shown, as these measured widths were likely wrong, because corresponding segments were extracted close to the tube ends or affected by contamination in the micrographs. The arrows at the bottom of the histogram indicate the diameters of helixes with different symmetries (see Fig. 5).

(B) 2D classification of 17,920 segments (2x binned, 1.65 Å/px) in cisTEM[1] detects tube diameter variation to a certain extent. Selected class averages for further processing are circled in red.

#### Extended Data Fig. 2: Fourier shell correlations (FSCs) for the helical and local reconstructions, respectively, at different stages of cryo-EM data processing.

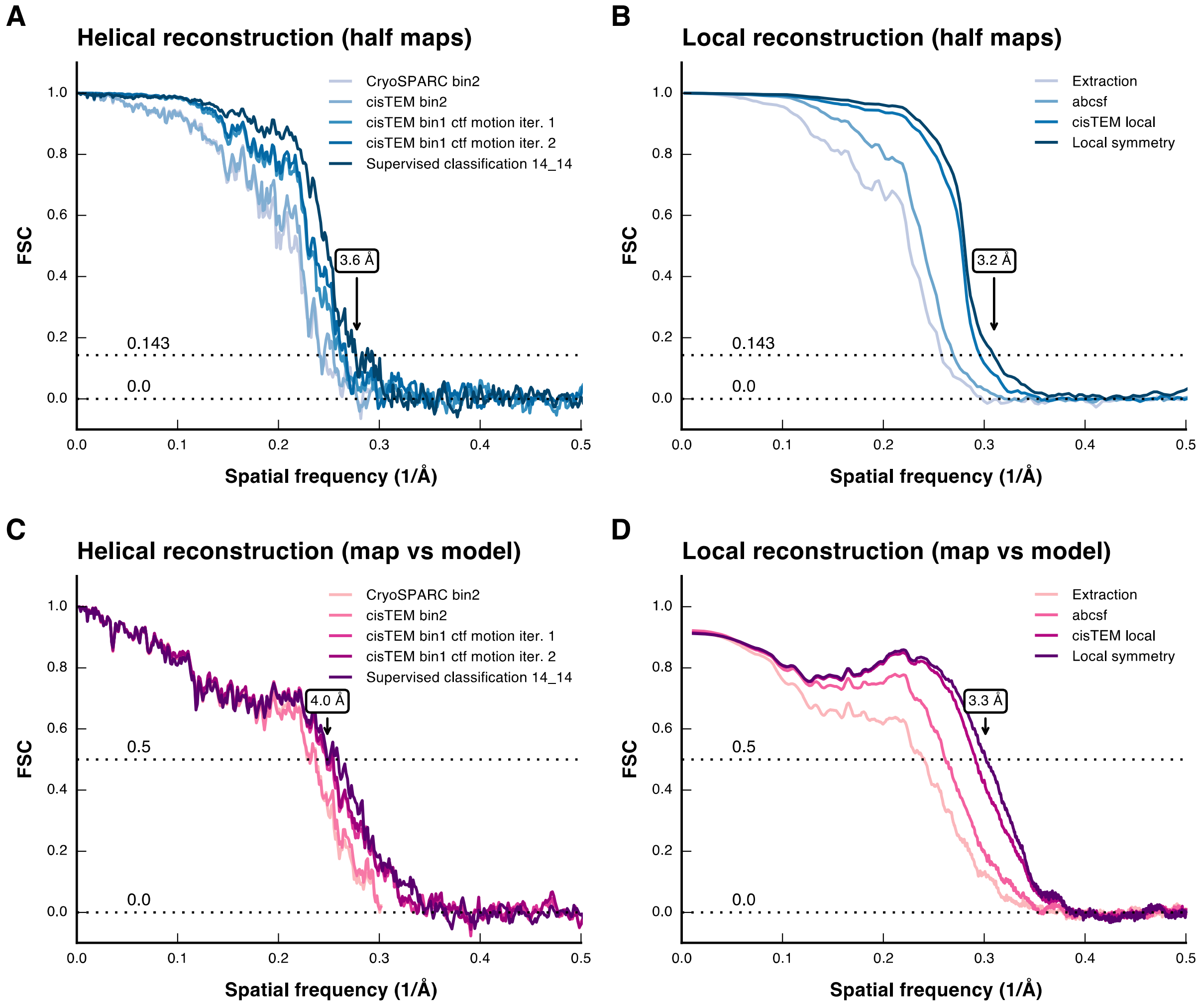


(A) FSC curves between half maps of the helical reconstruction. Correlation was calculated with e2proc3d.py from EMAN2[2] after masking the symmetrized half maps of the helical reconstructions. The final nominal resolution of 3.6 Å is indicated.

(B) FSC curves between half maps of the local reconstruction. Correlation was calculated with e2proc3d.py from EMAN2[2] after masking the half maps of the local reconstructions. The final nominal resolution of 3.2 Å is indicated.

(C) FSC curves between full maps and the final refined model of the helical reconstruction. The final refined model of a VP39 dimer was placed into the full map, rigid-body refined, and symmetry expanded to generate the full helix. We used the programs sfall and fftbig from CCP4[3] to calculate model structure factors and a model map. Correlation was then calculated with e2proc3d.py from EMAN2[2] after masking the symmetrized full map of the helical reconstructions and the model map, respectively. The final nominal resolution of 4.0 Å is indicated.

(D) FSC curves between full maps and the final refined model of the local reconstruction. Correlation was calculated with phenix.mtriage[4]. The final nominal resolution of 3.3 Å is indicated.

#### Extended Data Fig. 3: Depiction of local resolution of the cryo-EM map and B factors of the model

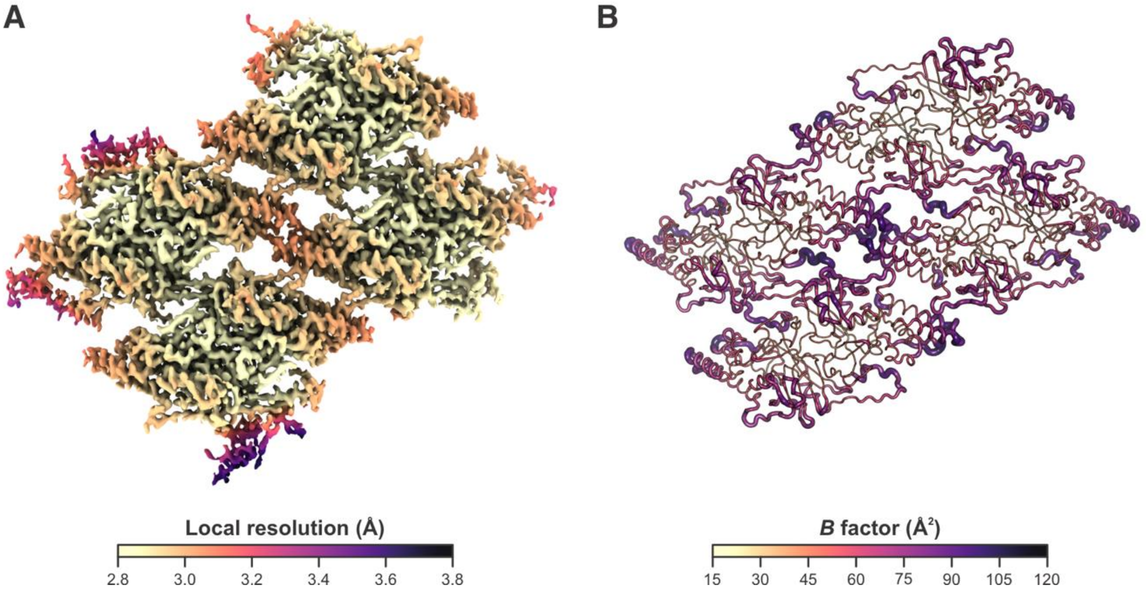


(A) Estimated local resolution mapped onto the local reconstruction of 4 adjacent dimers. Resolution estimates are color-coded from light (high resolution) to dark (low resolution). (B) Model of 4 adjacent dimers color coded by its refined B factors from light (low) to dark (high).

**Extended Data Fig. 4: Close-up view of the cryo-EM density map after local reconstruction with a nominal resolution of 3.2 Å.**

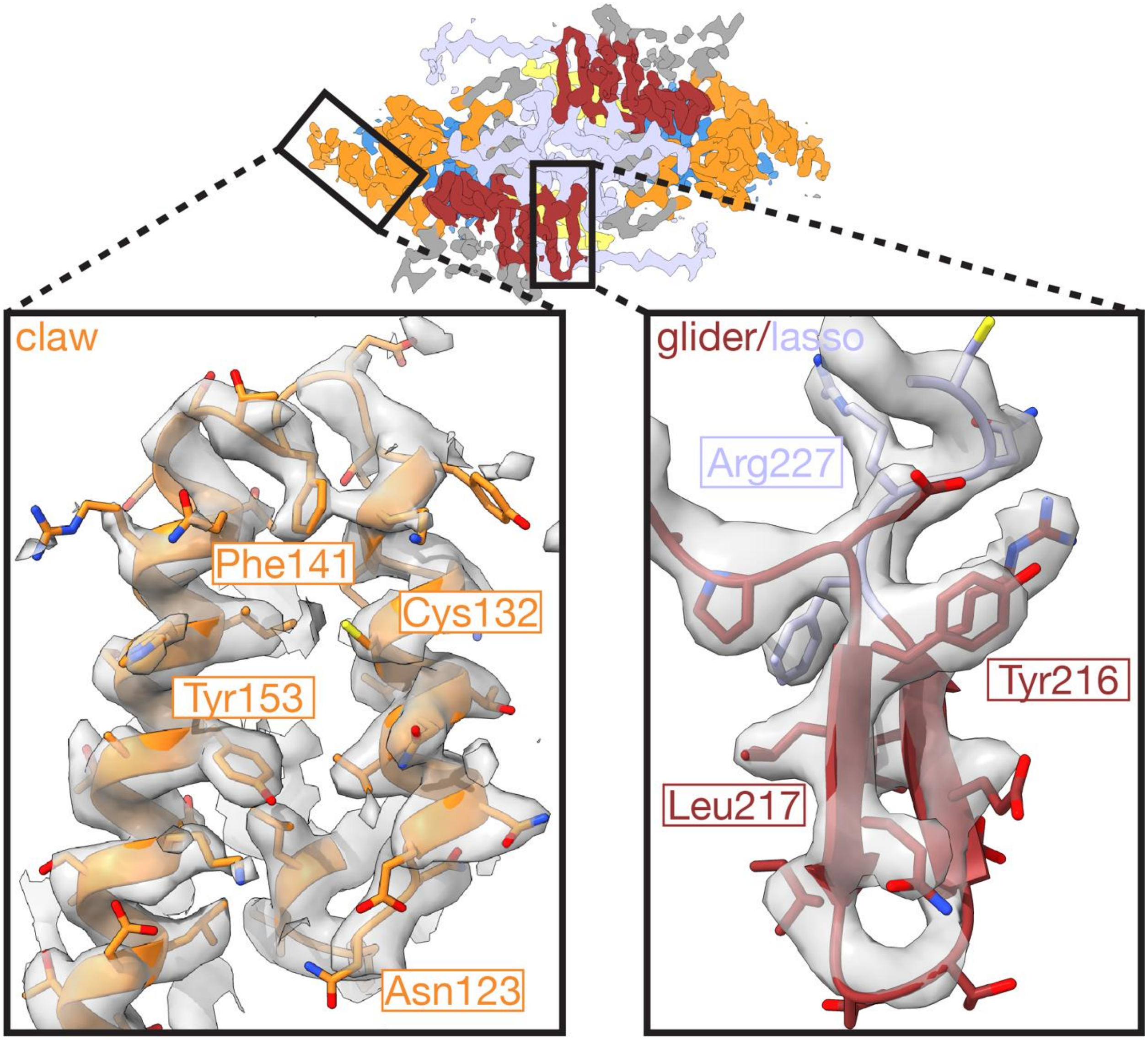


Residues 120-163 of the claw region (panel left; orange) and residues 213-229 of the glider and lasso region (panel right; red and light purple) are depicted.

**Extended Data Fig. 5: Zn-coordination site in the AcMNPV VP39 ZF region.**

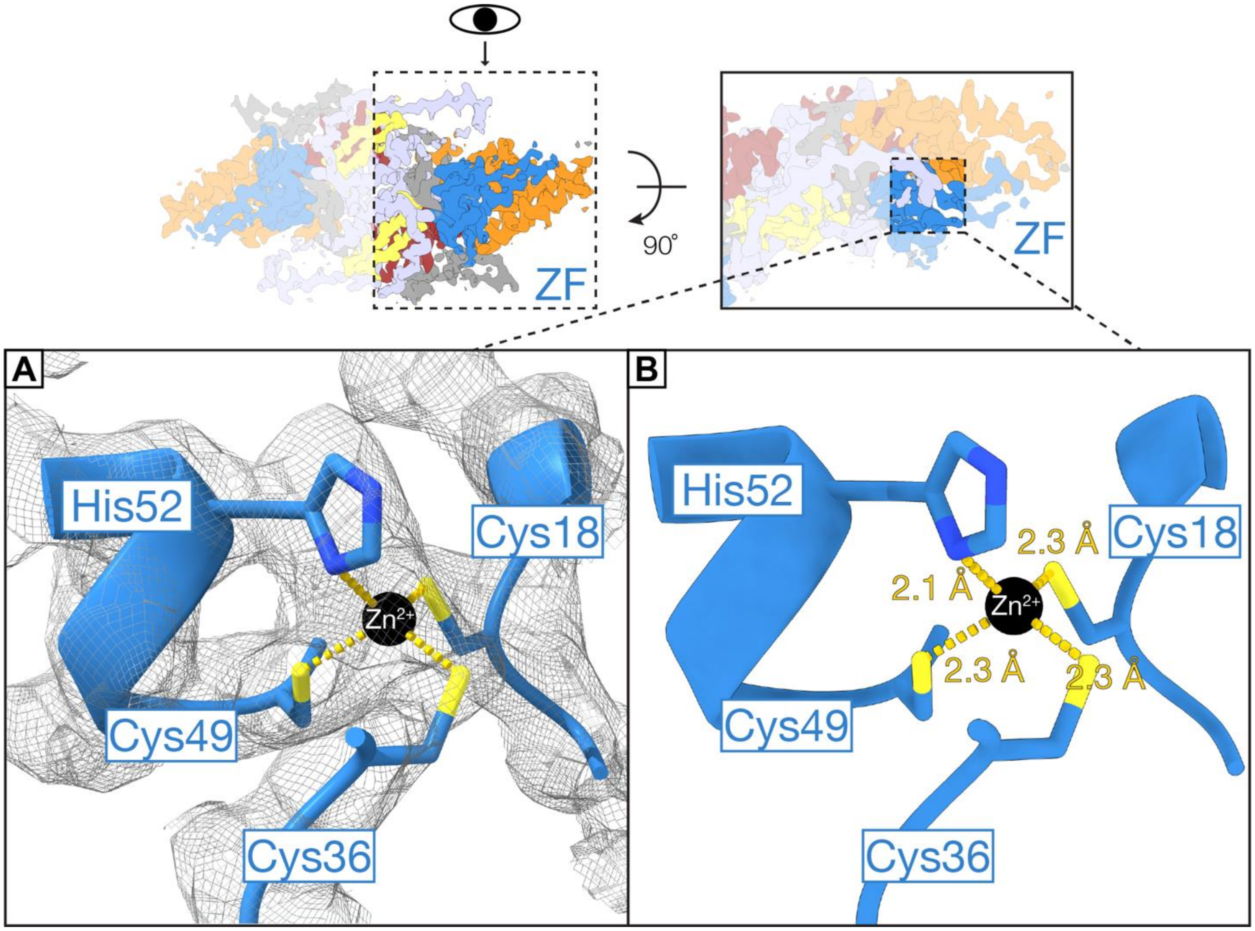


(A) Close-up view of the cryo-EM density at the Zn-coordination site. Cysteine 18, cysteine 36, cysteine 49 and histidine 52 coordinate a $Zn^{2+}$ ion (black sphere).

(B) Close-up view of the Zn-coordination site with nitrogen/sulphur atom-$Zn^{2+}$ distances shown. Distances were measured in ChimeraX[5].

**Extended Data Fig. 6: Electrostatic surface potential of the AcMNPV VP39 capsid.**

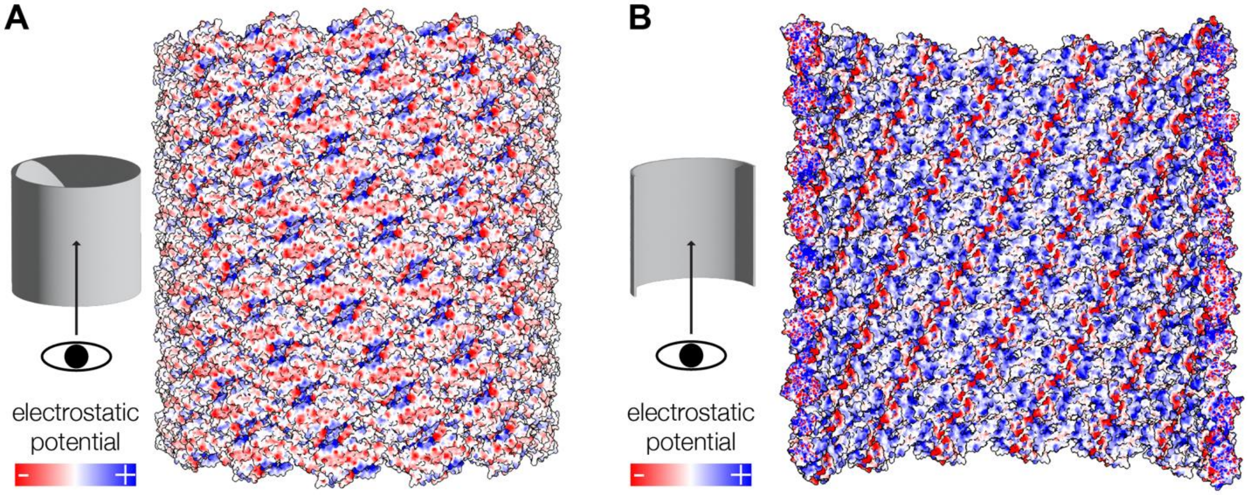


(A) Exterior view of the VP39 capsid. Surface representation of the VP39 model, which was fitted in the 3.6-Å cryo-EM capsid reconstruction. Surface is colored according to electrostatic potential from negative (red) to positive (blue).

(B) Luminal view of the VP39 capsid. Model fitting and electrostatic potential coloring were performed using ChimeraX[5].

**Extended Data Fig. 7: Two reduced cysteine pairs at the lateral inter-subunit interface.**

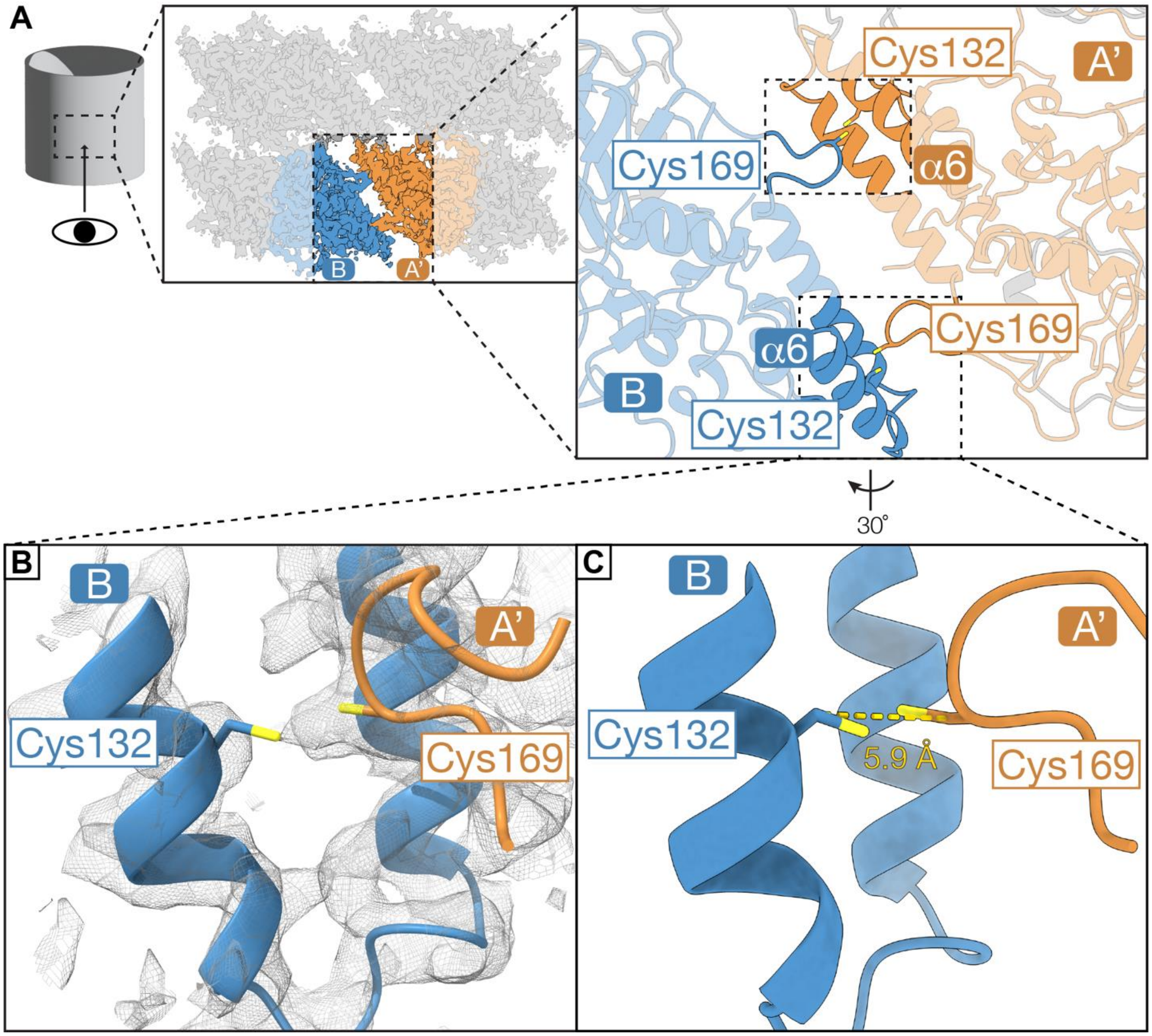


(A) Schematic overview of the location of the two potential disulfide pairs, viewed from the exterior side of the capsid. The four cysteines are located on the claw regions of two laterally adjacent subunits B and A'.

(B) Close-up view of the cryo-EM density around the two reduced cysteine residues cysteine 132 in monomer B and cysteine 169 in monomer A.

(C) Close-up view of one of the two candidate disulfide pairs with Cα-Cα distance shown. Distance was measured in ChimeraX[5].

**Extended Data Fig. 8: Electrostatic surface potential of baculoviral VP39 dimer predictions.**

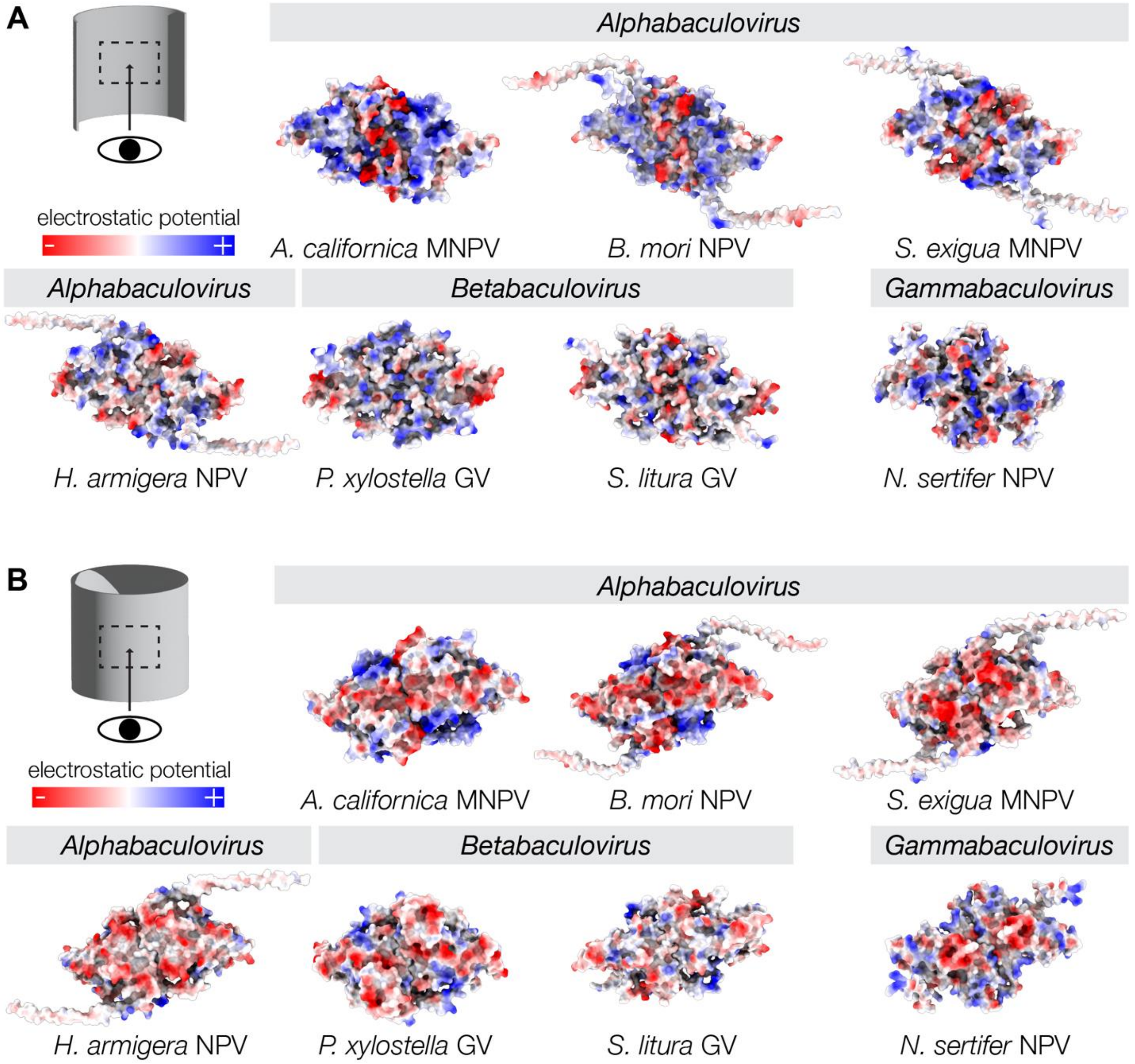


(A) Luminal view of VP39 dimer structure (AcMNPV) and AlphaFold2[6,7] predictions of the VP39 dimer. Electrostatic potential is colored from negative (red) to positive (blue).

(B) Exterior view of VP39 dimer structure and predictions with electrostatic potential colored accordingly.

MNPV: multiple nucleopolyhedrovirus; NPV: nucleopolyhedrovirus; GV: granulovirus.

**Extended Data Fig. 9: The conserved glycine 276 is located at the type-ii axial inter-subunit interface.**

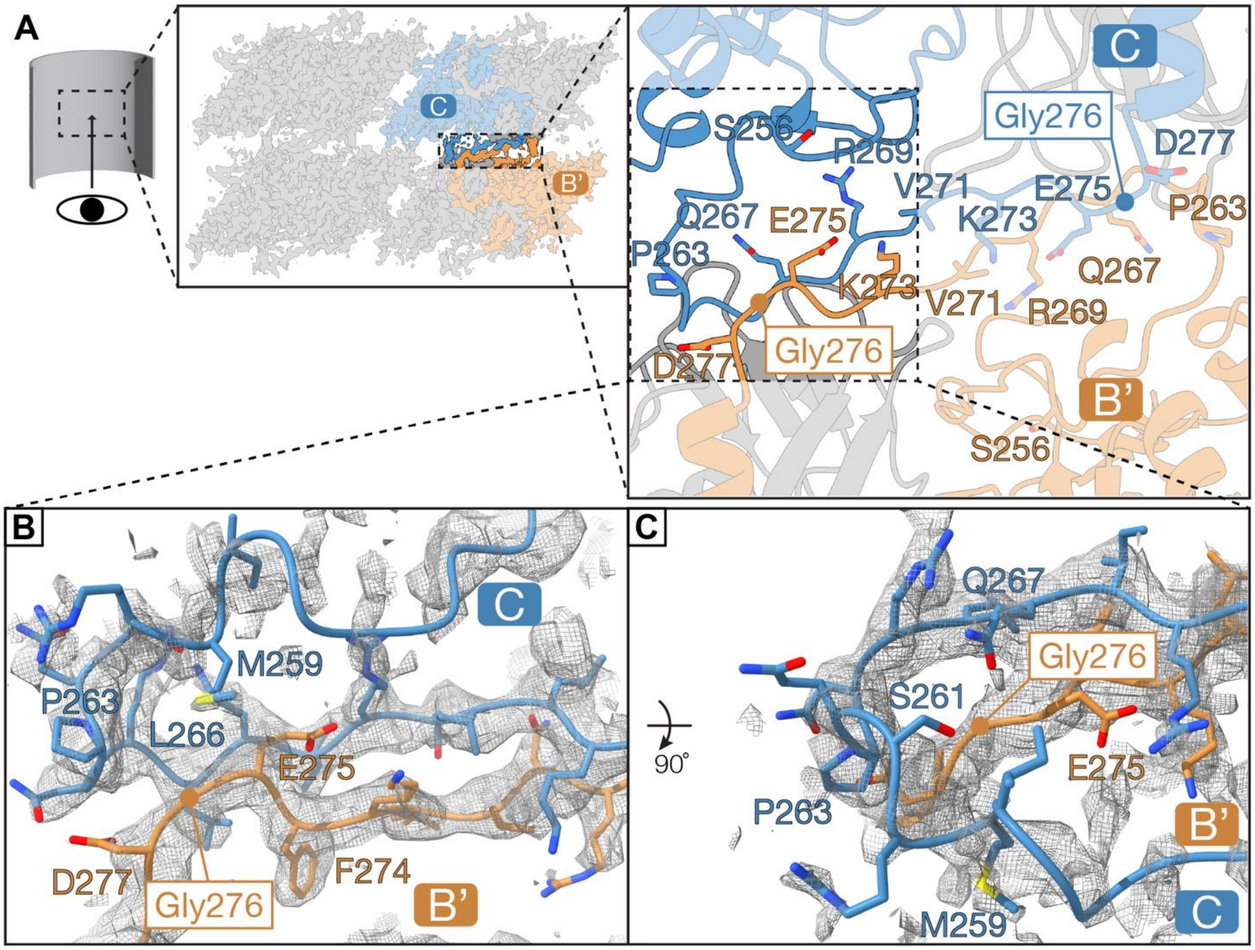


Morphologically aberrant nucleocapsids were observed in a mutagenesis screen with G276S[8].

(A) Schematic overview of the location of glycine 276, viewed from the luminal side of the capsid. Glycine 276 is nestled in between type-ii axial inter-subunit contacts on the lasso regions of two axially adjacent subunits B' and C. All interface residues are labeled.

(B) Close-up view of the cryo-EM density in the vicinity of glycine 276 in monomer B'.

(C) Top view of the immediate environment surrounding glycine 276 in monomer B'.

**Extended Data Fig. 10: Visualization of the candidate actin-binding residues 192-286.**

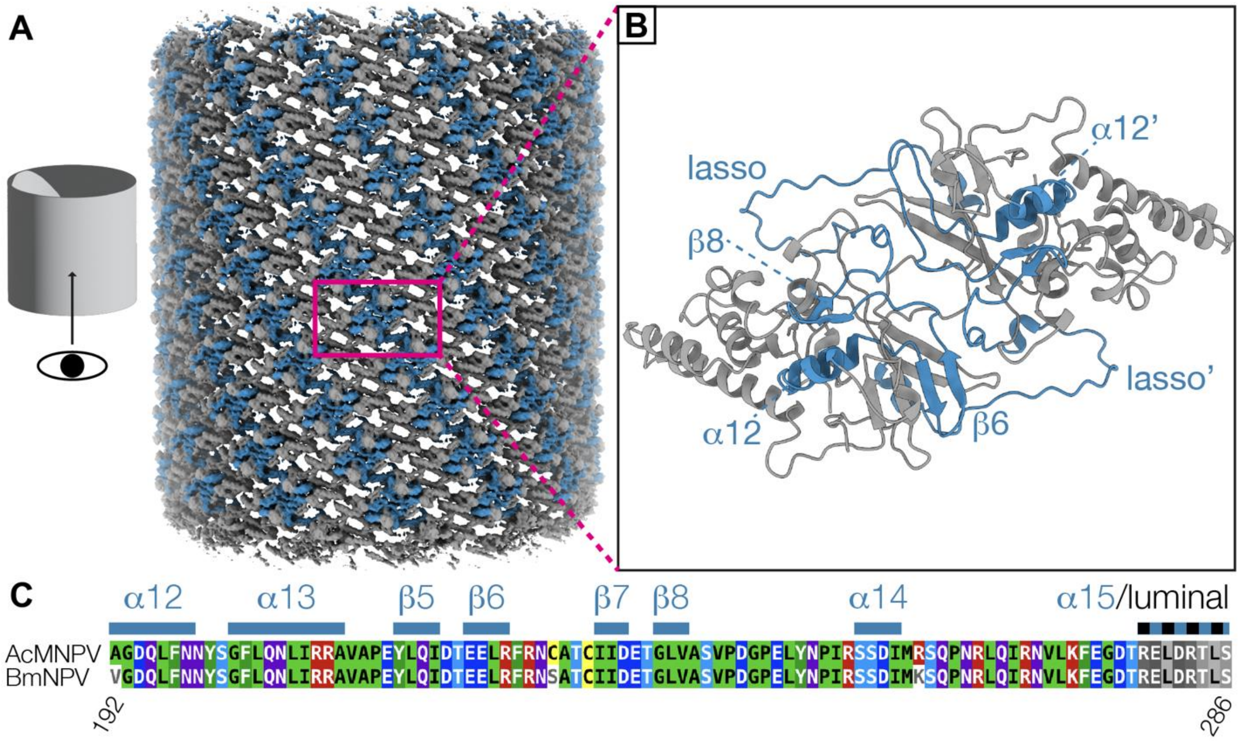


Residues 192-286 in BmNPV VP39 are required for nuclear actin polymerization[9].
(A) Cryo-EM map of the AcMNPV nucleocapsid with residues 192-286 colored in blue.
(B) Model of the AcMNPV dimer with residues 192-286 colored in blue.
(C) Sequence alignment of residues 192-286 of AcMNPV VP39 BmNPV VP39 with residues colored by identity (96.8% sequence identity). The last 8 residues (279-286; gray-scale coloring) are facing the luminal side of the capsid. Secondary structure elements as identified from our reconstruction are depicted above the sequence alignment. Sequence alignment was performed using MAFFT[10] and visualized with MView[11].

#### References


1. Grant, T., Rohou, A. & Grigorieff, N. cisTEM, user-friendly software for single-particle image processing. *eLife* **7**, e35383 (2018).
2. Tang, G. et al. EMAN2: An extensible image processing suite for electron microscopy. *Journal of Structural Biology* **157**, 38-46 (2007).
3. Winn, M.D. et al. Overview of the CCP4 suite and current developments. *Acta Crystallogr D Biol Crystallogr* **67**, 235-42 (2011).
4. Afonine, P.V. et al. New tools for the analysis and validation of cryo-EM maps and atomic models. *Acta Crystallographica Section D* **74**, 814-840 (2018).
5. Pettersen, E.F. et al. UCSF ChimeraX: Structure visualization for researchers, educators, and developers. *Protein Sci* **30**, 70-82 (2021).
6. Evans, R. et al. Protein complex prediction with AlphaFold-Multimer. *bioRxiv*, 2021.10.04.463034 (2022).
7. Jumper, J. et al. Highly accurate protein structure prediction with AlphaFold. *Nature* **596**, 583-589 (2021).
8. Katsuma, S. & Kokusho, R. A Conserved Glycine Residue Is Required for Proper Functioning of a Baculovirus VP39 Protein. *J Virol* **91**(2017).
9. Zhang, J., Li, Y., Zhao, S. & Wu, X. Identification of A functional region in Bombyx mori nucleopolyhedrovirus VP39 that is essential for nuclear actin polymerization. *Virology* **550**, 37-50 (2020).
10. Katoh, K., Rozewicki, J. & Yamada, K.D. MAFFT online service: multiple sequence alignment, interactive sequence choice and visualization. *Briefings in Bioinformatics* **20**, 1160-1166 (2017).
11. Madeira, F. et al. Search and sequence analysis tools services from EMBL-EBI in 2022. *Nucleic acids research* **50**, W276-W279 (2022).
